# Activation of the Maternal Genome Through Asymmetric Distribution of Oocyte-Genome-Associated Histone H3.3

**DOI:** 10.1101/2023.11.01.565208

**Authors:** Duancheng Wen, Zev Rosenwaks

## Abstract

Upon fertilization, the typically silent and epigenetically repressed oocyte genome undergoes activation, yet the precise mechanism remains unclear. The histone variant H3.3 is evenly distributed throughout the oocyte genome, suggesting its involvement in repression. This study reveals that oocyte-genome-associated H3.3 (oH3.3) undergoes asymmetric segregation among four-cell stage blastomeres, persisting in only two blastomeres through the blastocyst stage. These oH3.3-retaining blastomeres maintain a repressive state characterized by high levels of the chromatin marker H3K9me2. Intriguingly, single-cell RNA-seq analysis revealed asymmetric transcriptional activation between paternal and maternal genomes, with the maternal genome being considerably less active. We propose a model wherein oH3.3 and associated oocyte DNA co-segregate during mitosis, allowing two blastomeres to inherit oH3.3 and a strand of oocyte DNA from maternal chromatids. Meanwhile, the other blastomeres acquire newly synthesized DNA associated with the nascent histone H3, which lacks oocyte-specific repressive modifications. Consequently, full maternal genome activation occurs in two of the four-cell stage blastomeres, while the remaining two, which retain oH3.3, remain partially repressed. This study uncovers a previously unrecognized H3.3 mediated mechanism for maternal genome activation.

## Introduction

The genomes of sperm and oocytes possess distinct epigenetic features. The sperm genome is mainly packaged with protamines, while the oocyte genome is characterized by hypermethylation (1–4) and distinguished by nucleosomes that distribute the histone variant H3.3 uniformly across the entire genome (5). Upon fertilization, the previously transcriptionally inactive sperm and oocyte genomes unite to create a diploid zygotic genome, which then experiences comprehensive epigenetic reprogramming, eventually leading to zygotic genome activation (ZGA) (6). Numerous studies point towards divergent reprogramming dynamics in paternal and maternal genomes. Specifically, demethylation in the maternal genome tends to be passive and DNA replication-dependent, whereas the paternal genome undergoes active DNA demethylation during the early phases of embryonic development (7–12). Additionally, the paternal genome generally exhibits higher transcriptional activity than the maternal genome (13) and the activation of the maternal genome tends to lag behind that of the paternal genome during ZGA (6, 14). A recent study demonstrated that, during embryogenesis, the maternal genome segregates independently from the paternal genome via a dual-spindle formation mechanism (15); however, the biological implications of this independent segregation remain unclear. Collectively, it is widely accepted that the activation of the maternal genome follows a fundamentally different mechanism compared to the activation of the paternal genome, which takes place as early as a few hours after fertilization in mice.

The histone variant H3.3 in mammals is encoded by two distinct genes, *h3f3a* (H3.3A) and *h3f3b* (H3.3B), both of which produce an identical protein product (16, 17). H3.3 is constitutively expressed in cells and incorporated into chromatin via a DNA synthesis-independent pathway during and outside the S phase (18). The study of H3.3 has garnered increasing interest in the field of developmental biology due to its unique roles in remodeling both male and female genomes during fertilization and early embryonic development. H3.3A and H3.3B are expressed during oogenesis and throughout preimplantation stages and can functionally compensate for each other (19). This functional compensation is evidenced by the observation that the deletion of a single gene does not result in any noticeable developmental phenotype (20, 21).

H3.3 is known to be enriched in the genomes of both sperm (sH3.3) and oocytes (oH3.3) (5, 22). The oocyte cytoplasm is also stockpiled with maternal H3.3 mRNAs (mH3.3) and zygotic expression of H3.3 (zH3.3) begins as early as the two-cell stage (22). We previously reported that sH3.3 is rapidly removed from the sperm genome post fertilization (23), and mH3.3-produced protein is deposited into the paternal genome as early as 1 hour after fertilization, remaining detectable until the morula stage (23). Our research (19, 23), along with that of others (5, 21, 24), has demonstrated that mH3.3 is essential for fertilization and early embryonic development, emphasizing its importance in regulating gene expression and reprogramming both the paternal and maternal genomes. However, due to the coexistence of oH3.3 with mH3.3 and zH3.3 in early embryos, tracking the specific dynamics of these different sources of H3.3 has proven technically challenging. As a result, the fate of oH3.3 during early embryonic development and its potential role in epigenetic regulation of the maternal genome after fertilization remains unknown.

In this study, using H3.3B-HA reporter mice, we discovered that oH3.3 is asymmetrically segregated and distributed among the four-cell stage blastomeres, with only two blastomeres retaining oH3.3 exclusively throughout the preimplantation stages. These oH3.3-retaining cells maintain a repressive chromatin state, characterized by high levels of the repressive histone H3 modification, H3K9me2. They are located within the trophoblast layer rather than contributing to the inner cell mass (ICM) of a blastocyst. We also demonstrated that full maternal genome activation is delayed until the four-cell stage.

## Results

### oH3.3B-HA Retained in Maternal Nuclei up to the Two-Cell Stage

We introduced a small C-terminal HA tag fused to the final coding exon of the H3.3B gene and generated H3.3B-HA reporter mice (22). This tag was accompanied by an internal ribosomal entry site (IRES) and a separately translated EYFP or mCherry reporter, which allowed us to study the expression and localization of the H3.3B gene during development (**Fig. 1A. 1B**) (22).

**Figure 1.**
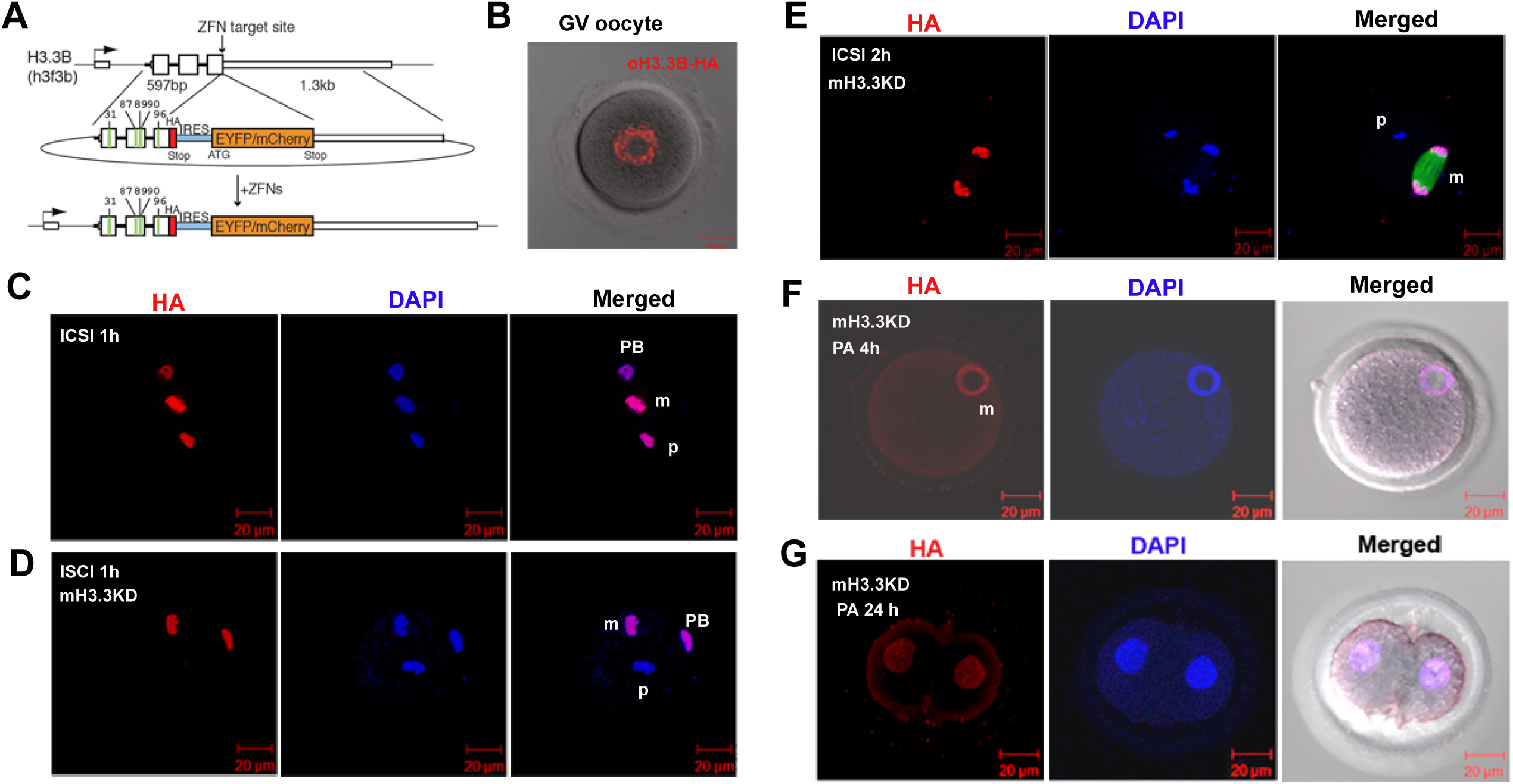
Retention of oH3.3 in the maternal genome post-fertilization. (**A**) Schematic representation of the H3.3B-HA-IRES-EYFP/mCherry constructs targeted by zinc finger nucleases. (**B**) All oocytes from heterozygous H3.3B-HA-tagged females display HA-positive staining in the nucleus, irrespective of oocyte genotype (representative GV oocyte shown). (**C**) Deposition of mH3.3 to the paternal genome (p) 1-hour post intracytoplasmic sperm injection (ICSI) with wild-type sperm. PB: polar body. m: maternal genome. (**D**) Efficient knockdown of mH3.3 achieved by injecting siRNAs (targeting both *h3f3a* and *h3f3b*) into oocytes prior to ICSI, preventing mH3.3 deposition to the paternal genome (p), while oH3.3 is detected in the maternal genome (m) and PB. (**E**) The mH3.3-knockdown oocyte is fertilized with a wild-type sperm. No mH3.3 deposition is observed on the paternal genome 2 hours post-ICSI, but a strong oH3.3 (HA) signal is detected in the maternal genome (m) with spindle formation (Green: α-tubulin). (**F**) A parthenogenetically activated (PA) oocyte with mH3.3KD 4 hours after activation displays oH3.3B-HA positive staining in the maternal pronucleus (m). (**G**) A parthenogenetically activated 2-cell embryo (PA 24h) with mH3.3KD exhibits oH3.3B-HA positive staining in the nuclei of both blastomeres.

To detect the oH3.3 protein (oH3.3B-HA) during fertilization and early embryogenesis, it is essential to eliminate signals originating from maternally stored mH3.3B-HA mRNA within the oocyte cytoplasm. To achieve this, we microinjected small interfering RNAs (siRNAs) specifically targeting H3.3A and H3.3B (siH3.3) into MII oocytes (19). Afterward, the injected oocytes underwent fertilization via intracytoplasmic sperm injection (ICSI) using wild-type sperm. In oocytes without siRNA injections, mH3.3B-HA was detected on both the sperm (mH3.3) and oocyte (oH3.3+mH3.3) genomes one hour (1h) post-ICSI using an HA antibody (**Fig. 1C**). In contrast, siH3.3 knockdown (KD) oocytes exhibited no mH3.3B-HA staining on the sperm genome, with only HA positive staining in the oocyte genome (oH3.3) in the embryos 1h and 2h post-ICSI (**Fig. 1D, 1E**), indicating the efficient depletion of mH3.3 mRNAs by siH3.3. Since male and female genomes undergo fusion starting from 4h after fertilization, we used parthenogenetically activated (PA) oocytes to continue monitoring oH3.3 in the mH3.3 KD embryos. This HA staining persisted in the female pronucleus (4h) (**Fig. 1F**) and in the nuclei of the 2-cell stage embryos (24h) (**Fig. 1G**) in siH3.3 KD PA oocytes, suggesting that oH3.3B-HA remains present in the maternal genome after fertilization/PA activation and is retained up to the 2-cell stage.

### Asymmetric Distribution of oH3.3B-HA in Diploid PA Embryos: Four Blastomeres Retained oH3.3

Since mH3.3 KD significantly affects the developmental potency of embryos (19, 23, 24), these embryos are unsuitable for further tracking of oH3.3 during embryogenesis. To eliminate the mH3.3B-HA mRNA in the oocyte cytoplasm, we performed spindle exchange between H3.3B-HA oocytes obtained from heterozygous H3.3B-HA females and WT oocytes (B6D2F1), creating an oocyte with an H3.3B-HA nucleus and WT ooplasm (**Fig. 2A**). Oocytes from heterozygous H3.3B-HA mice produce two genotypes, e.g., H3.3B-HA^pos^ (bearing the HA allele) and H3.3B-HA^neg^ (carrying the WT allele) oocytes (**Fig. 2A**). However, oH3.3B-HA is deposited onto the oocyte genome regardless of its genotype during oogenesis, as evidenced by all the oocytes from heterozygous H3.3B-HA female mice being HA positive in the genome (**Fig. 1B**). Moreover, these two genotypes can be distinguished by the expression of EYFP after the 2-cell stage (22). Utilizing these reconstructed oocytes allows us to specifically monitor the fate of the oH3.3B-HA protein inherited from the oocyte genome during early embryogenesis using an HA antibody (**Fig. 2B**).

**Figure 2.**
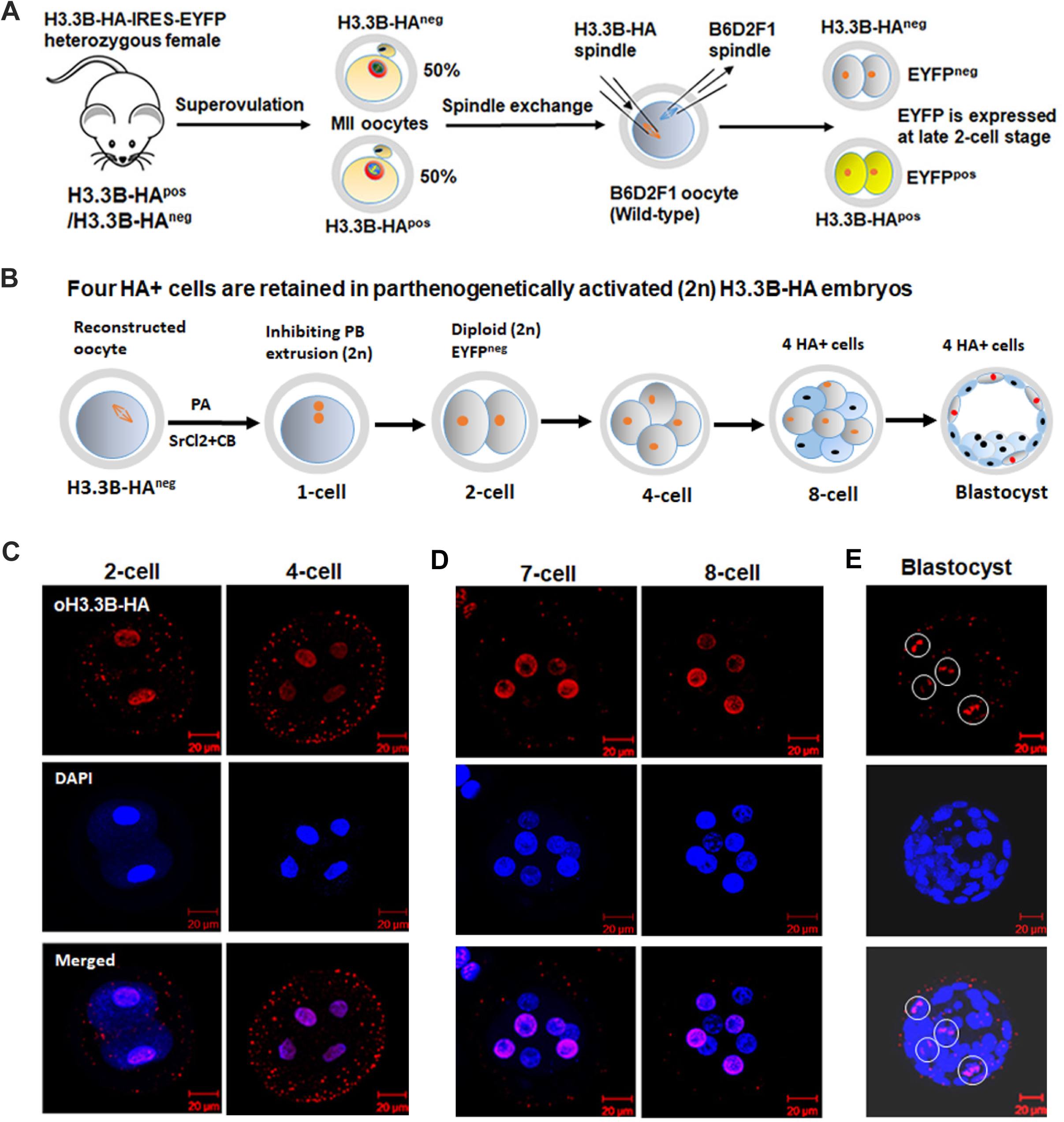
Asymmetric segregation and distribution of oH3.3 in parthenogenetically activated embryos. (**A**-**B**) Schematic illustration of tracking oH3.3 in diploid parthenogenetic embryos. H3.3B-HA heterozygous females produce oocytes with two genotypes: H3.3B-HA-positive (pos) and H3.3B-HA-negative (neg), which represent the wild-type allele. The genome of oocytes with both genotypes is enriched with oH3.3B-HA, which can be detected using HA-specific antibodies. Maternal H3.3B-HA (mH3.3) is eliminated through spindle exchange with a wild-type oocyte (**A**). Approximately 50% of the reconstructed oocytes are of the H3.3B-HA^neg^ genotype and can be distinguished by the expression of EYFP after the 2-cell stage (only H3.3B-HA^pos^ oocytes express EYFP). The reconstructed oocytes are parthenogenetically activated with SrCl2 to generate diploid embryos by inhibiting polar body extrusion using cytochalasin B (CB). The oH3.3-HA is tracked by HA staining in preimplantation embryos. (**C**) The oH3.3B-HA can be detected in the genome of every blastomere at the 2-cell (n=4) and 4-cell (n=4) stage embryo. (**D**-**E**) Only four blastomeres in the nucleus exhibit oH3.3B-HA positive staining in 7-cell/8-cell (n=7) (D), and blastocyst (n=5) (E) stage embryos.

The reconstructed H3.3B-HA oocytes were parthenogenetically activated with SrCl2 and supplemented with cytochalasin B (CB) to inhibit polar body (PB) extrusion, thus creating a diploid karyotype (**Fig. 2B**). Embryos negative for EYFP (and consequently, negative for zygotic expression of H3.3B-HA) were harvested for HA staining and imaging. As anticipated, all the blastomeres from 2-cell and 4-cell stage embryos exhibited positive HA staining in the nucleus (**Fig. 2C**). This result suggests that oH3.3B-HA is retained not only within the genome but also distributed evenly among all blastomeres up to the 4-cell stage in the diploid parthenogenetically activated (2n PA) embryos.

Intriguingly, our observations revealed a shift in the later stages of embryonic development. Starting from the 8-cell stage and continuing through to the blastocyst stage, only four blastomeres in the 2n PA embryos exhibited positive H3.3B-HA (oH3.3B-HA) staining (**Fig. 2D, 2E**). The intensity of the oH3.3B-HA staining remained relatively consistent in the nuclei throughout various developmental stages, contrasting with the rapidly decreasing intensity of mH3.3B-HA staining that becomes undetectable in morulae and blastocysts (19) as embryonic development progresses. This observation implies that oH3.3B-HA is not diluted or removed after the 4-cell stage, persisting until the blastocyst stage while being confined to four blastomeres within the 2n PA embryos.

This intriguing finding suggests that following the third embryonic cleavage, the segregation and distribution of oH3.3B-HA among blastomeres in 2n PA embryos exhibit an asymmetric pattern. Given that 2n PA embryos contain two sets of maternal chromosomes and each chromosome consists of a double-stranded DNAs (Watson (W-strand) and Crick (C-strand) strands), each blastomere at the 4-cell stage thus can receive a complete set of chromatids comprising one nascent DNA strand and one oDNA strand (W-oDNA:: C-nascent DNA and W-nascent DNA:: C-oDNA). The observation that only four cells consistently display oH3.3B-HA positive staining in the embryos from the 4-cell stage to the blastocyst stage suggests that oH3.3B-HA-associated chromatids are not randomly distributed among the blastomeres during embryogenesis. Instead, a specific allocation process occurs, wherein one set of chromatids containing oH3.3B-HA/oDNAs is distributed together into one daughter cell. This unique allocation results in a maximum of only four cells exhibiting oH3.3B-HA positive staining in the 2n PA embryos across all preimplantation stages after the 4-cell stage.

### Asymmetric Distribution of oH3.3B-HA in Fertilized Embryos: Two Blastomeres Retain oH3.3, and Potential Loss during Embryo Development

To determine whether this asymmetric pattern of oH3.3B-HA distribution also occurs in normal fertilized embryos, we investigated the oH3.3B-HA pattern in fertilized embryos produced through ICSI using WT sperm. In these embryos, we anticipated observing oH3.3B-HA positive staining in only two cells throughout the preimplantation stages, as they contain just one set of oocyte chromosomes. Considering that the reconstructed H3.3B-HA oocytes cannot withstand further micromanipulation with ICSI and that mH3.3B-HA diminishes by the morula stage, we collected H3.3B-HA embryos at the 8-cell to blastocyst stages produced by ICSI with WT sperm for HA staining and imaging. Only EYFP-negative (H3.3B-HA negative) embryos under a fluorescence microscope were selected for further analysis (**Fig. 3A**). Indeed, we identified only two HA-positive staining cells in the 8-cell stage embryos (**Fig. 3B**) and a maximum of two HA-positive cells in all the blastocysts we observed (**Fig. 3C, 3D**). Our results confirm that oH3.3B-HA is also asymmetrically segregated and distributed in fertilized embryos. Only two blastomeres retain oH3.3B-HA throughout the preimplantation stage, aligning with the presence of one set of oocyte chromosomes in the fertilized embryo. The cell containing oH3.3B-HA at the metaphase stage showed a mix of chromosomes. Some chromosomes were positively stained with oH3.3B-HA (oDNAs), whereas others (paternal and nascent maternal DNAs) were not stained (**Fig. 3E, 3F**). This provides strong evidence that all oocyte genome chromatids associated with oH3.3B-HA segregate and are collectively distributed into a single cell. Notably, all the oH3.3B-HA positive cells were located in the trophoblast layer, with no cells showing oH3.3B-HA positive staining identified in the inner cell mass (ICM) of the fertilized blastocysts (**Fig. 3C-, 3G-3I**). Interestingly, our analysis of morulae and blastocysts revealed that most embryos exhibited only one HA-positive cell (20 out of 25 embryos) (**Fig. 3G-3I**). This observation suggests the potential loss or elimination of these specific cells during the developmental process.

**Figure 3.**
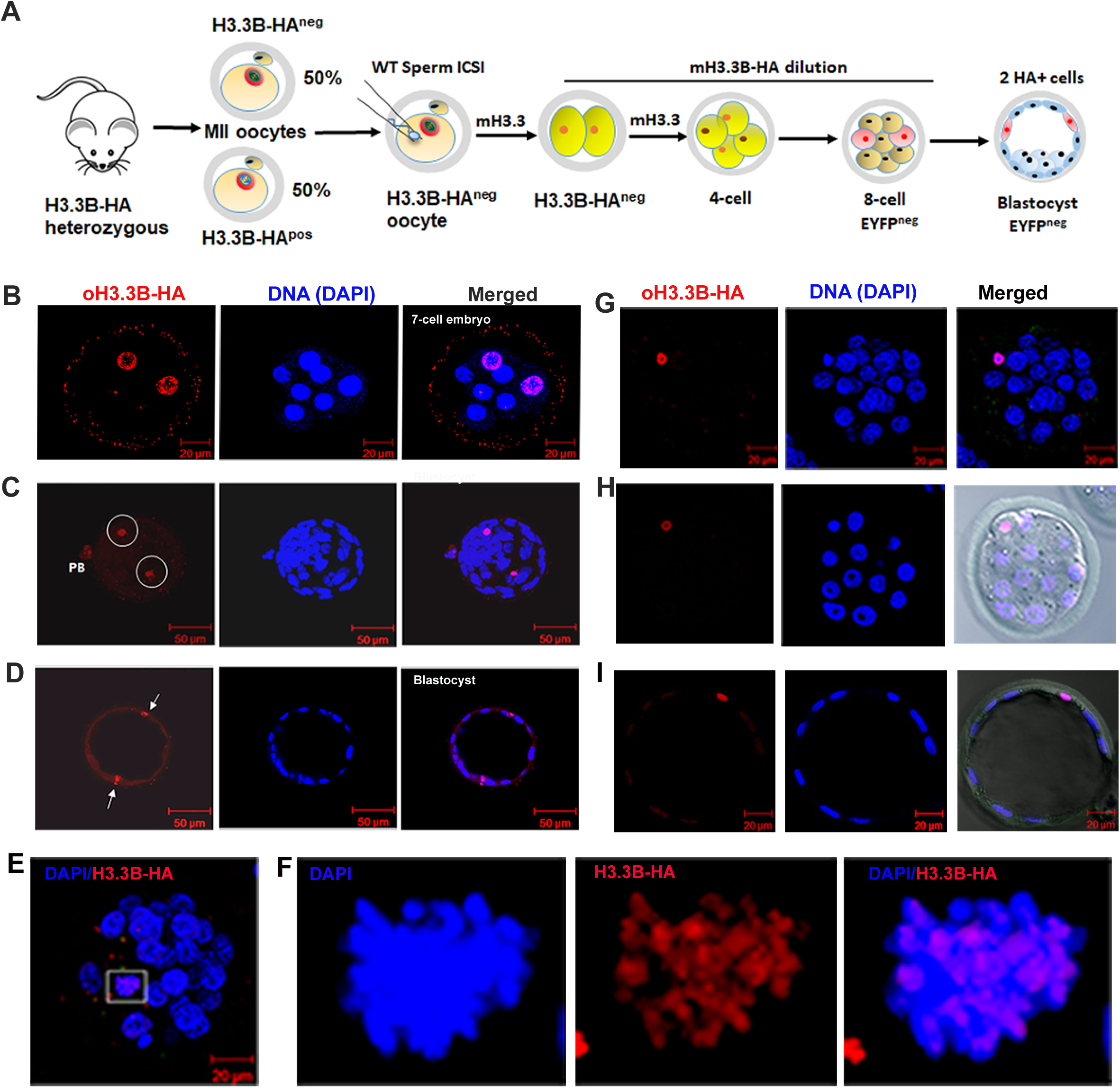
oH3.3 retention in fertilized preimplantation embryos. (**A**) Schematic illustration depicting the detection of oH3.3B-HA in fertilized embryos. Oocytes obtained from H3.3B-HA heterozygous females consist of 50% H3.3B-HA^pos^ and H3.3B-HA^neg^ genotypes, both containing mH3.3B-HA (maternal H3.3B-HA mRNA). These oocytes are fertilized with wild-type sperm via ICSI. The mH3.3B-HA is diluted by the morula stage and is undetectable at the blastocyst stage, as evidenced by the absence of EYFP fluorescence, while oH3.3B-HA continues to be detectable in a maximum of two cells by HA staining in the blastocyst. (**B**) Two blastomeres in the 7-cell embryo (n=1) show oH3.3B-HA positive staining in the genome. (**C**) A full projection of a fertilized blastocyst reveals two cells with oH3.3B-HA positive staining (n=5). PB: the 2nd polar body. (**D**) A single plane image of a fertilized blastocyst displays two oH3.3B-HA positive cells localized in the trophoblast layer. (**E**-**F**) An early fertilized blastocyst with the oH3.3 retaining cell in the metaphase stage demonstrates the intermingling of chromosomes. Some chromosomes are positively stained with oH3.3B-HA (oDNAs), while others (paternal and nascent maternal DNAs) are not stained. (**F**) Digital magnification of the oH3.3B-HA retaining cell shown in (**E**). (**G**-**I**) A single cell exhibits oH3.3B-HA positive staining in morula and blastocysts (n=20) from three independent experiments. One blastomere with HA+ possesses a more compact nucleus than other cells in a morula (**G**) and an early blastocyst (**H**). However, the size of the blastomere containing oH3.3 is similar to other blastomeres, ruling out the possibility of it being a polar body (**H**). A single plane image of a blastocyst highlights the HA+ cell’s location within the trophoblast layer (**I**).

### oH3.3-Retaining Cells Exhibit Compact Nuclei and Staining with Repressive H3K9me2 Modification

We typically observed a loss of H3.3B-HA-retaining cells in blastocysts and noticed that these cells exhibited nuclear morphological differences. Compared to other blastomeres, many of these oH3.3B-HA retaining cells appeared to have more compact nuclei, specifically in the blastocyst stage embryos (**Fig. 3G-3I**). However, the size of the blastomere containing oH3.3 is comparable to that of other blastomeres, ruling out the possibility of it being a polar body (**Fig. 3H**). The oH3.3-retaining cells display more compact nuclei than other blastomeres, suggesting a potential association with a repressive chromatin state. To further investigate this, we stained fertilized embryos using antibodies against H3 dimethyl lysine 9 (H3K9me2), a widely recognized marker for constitutive heterochromatin (25–27). Remarkably, the oH3.3B-HA-positive cells with compact nuclei displayed prominent staining for H3K9me2, providing strong evidence of their association with a repressed heterochromatin state (**Fig. 4A, 4B**).

**Figure 4.**
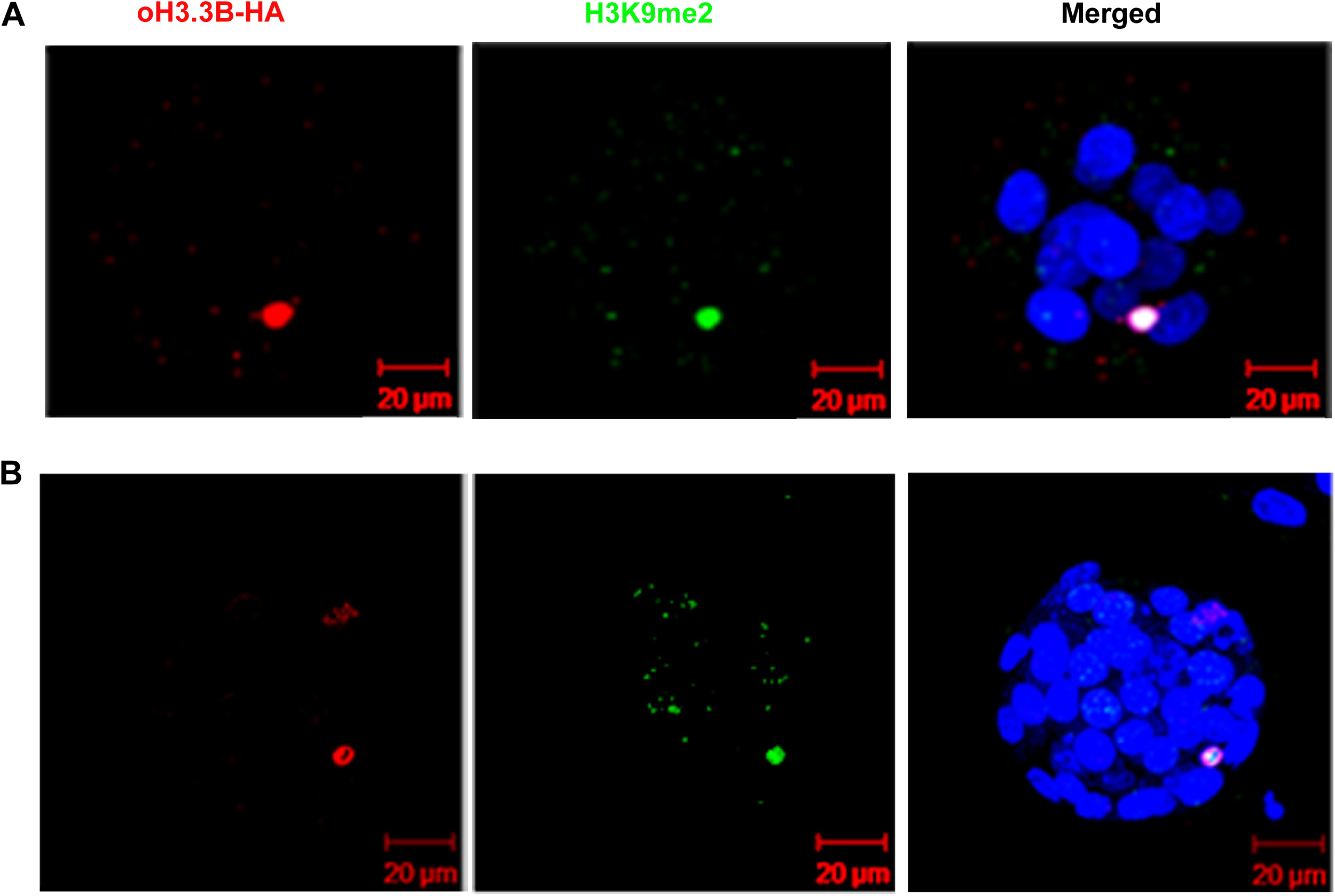
oH3.3-retaining cells demonstrate high levels of the repressive chromatin marker, H3K9me2 modification. (**A**) A cell in the morula stage, retaining oH3.3, displays pronounced staining for both H3K9me2 and H3.3B-HA (n=1). (**B**) Co-staining of H3.3B-HA and H3K9me2 in a blastocyst (n=3).

### Asymmetric Transcriptional Activation of Paternal and Maternal Genomes

We examined publicly available single-cell RNA-seq data from individual blastomeres of preimplantation mouse embryos (32). We found that paternal H3.3B starts expressing in one of the early 2-cell stage blastomeres (**Fig. 5A**). This observation was further confirmed through IF staining using H3.3B-HA in 2-cell embryos (**Fig. 5B**). While the average expression level of maternal H3.3B decreases from the zygote to the 8-cell stage, the level of paternal H3.3B shows an increasing trend from the 2-cell to the 8-cell stage (**Fig. 5C**). We conclude that paternal H3.3B activation begins at the early 2-cell stage, while the activation of the maternal H3.3B gene is deferred until the 4-cell stage. Intriguingly, one or two blastomeres in each 8-cell stage embryo exhibit significant depletion of the maternal H3.3B level. Specifically, five cells display levels at 1%-25% of the average in three of the embryos, while one of the seven blastomeres shows a level at 47% of the average in an 8-cell embryo (**Fig. 5D, 5E**). This pattern is not observed in paternal H3.3B (**Fig. 5D, 5E**). These observations indicate that the H3.3B gene from the maternal allele might be suppressed in one or two blastomeres within the embryo. This aligns with the presence of one or two oH3.3-retaining cells observed in preimplantation embryos. Thus, we consider these "H3.3B-low cells" to be oH3.3-retaining cells.

**Figure 5.**
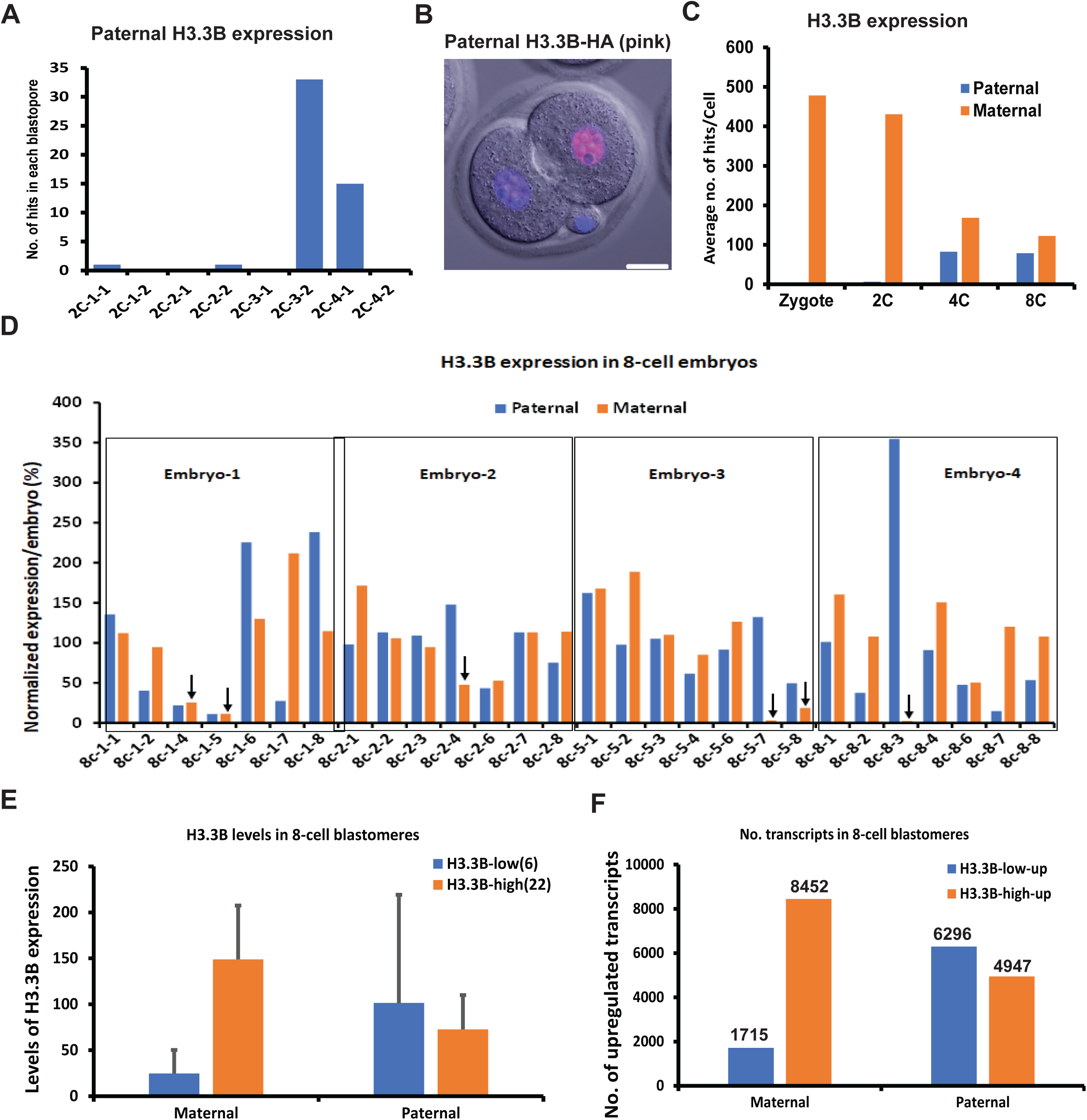
Asymmetric transcriptional activation of paternal and maternal genomes. (**A**) Activation of paternal H3.3B (h3f3b) occurs in one of the early 2-cell blastomeres. (**B**) Immunofluorescence (IF) staining reveals the presence of paternal H3.3B-HA in one of the early 2-cell stage blastomeres (pink color). Scale bar: 20 μm. (**C**) Comparison of average levels of H3.3B from paternal and maternal genomes in early-stage embryos. (**D** Expression patterns of H3.3B within individual blastomeres of 8-cell stage embryos. One or two blastomeres display a marked reduction of maternal H3.3B (indicated by arrow). (E) Average levels of H3.3B in the H3.3B-low (6 cells) and H3.3B-high (22 cells) 8-cell stage blastomeres. (F) No. of upregulated transcripts in the H3.3B-low cells and H3.3B-high cells in 8-cell embryos.

Despite the technical challenges of distinguishing oocyte-stored transcripts from de novo transcripts originating from the maternal genome post-fertilization, we hypothesize that cells retaining oH3.3, with their maternal genome in a repressive state, might influence global de novo transcription. To this end, we examined the number of upregulated transcripts from the maternal genome at the 8-cell stage by comparing the average level of each transcript at the 8-cell stage to that at the 4-cell stage. Notably, the upregulated transcripts identified from the maternal genome are significantly fewer in the H3.3B-low cells than in the H3.3B-high cells (**Fig. 6F**). Meanwhile, the number of upregulated transcripts from the paternal genome in both H3.3B-low and H3.3B-high cells is not significantly different (**Fig. 6F**). This suggests that the maternal genome with oH3.3 is less transcriptionally active than both the paternal genome and the maternal genome without oH3.3 during the early embryonic stages.

**Figure 6.**
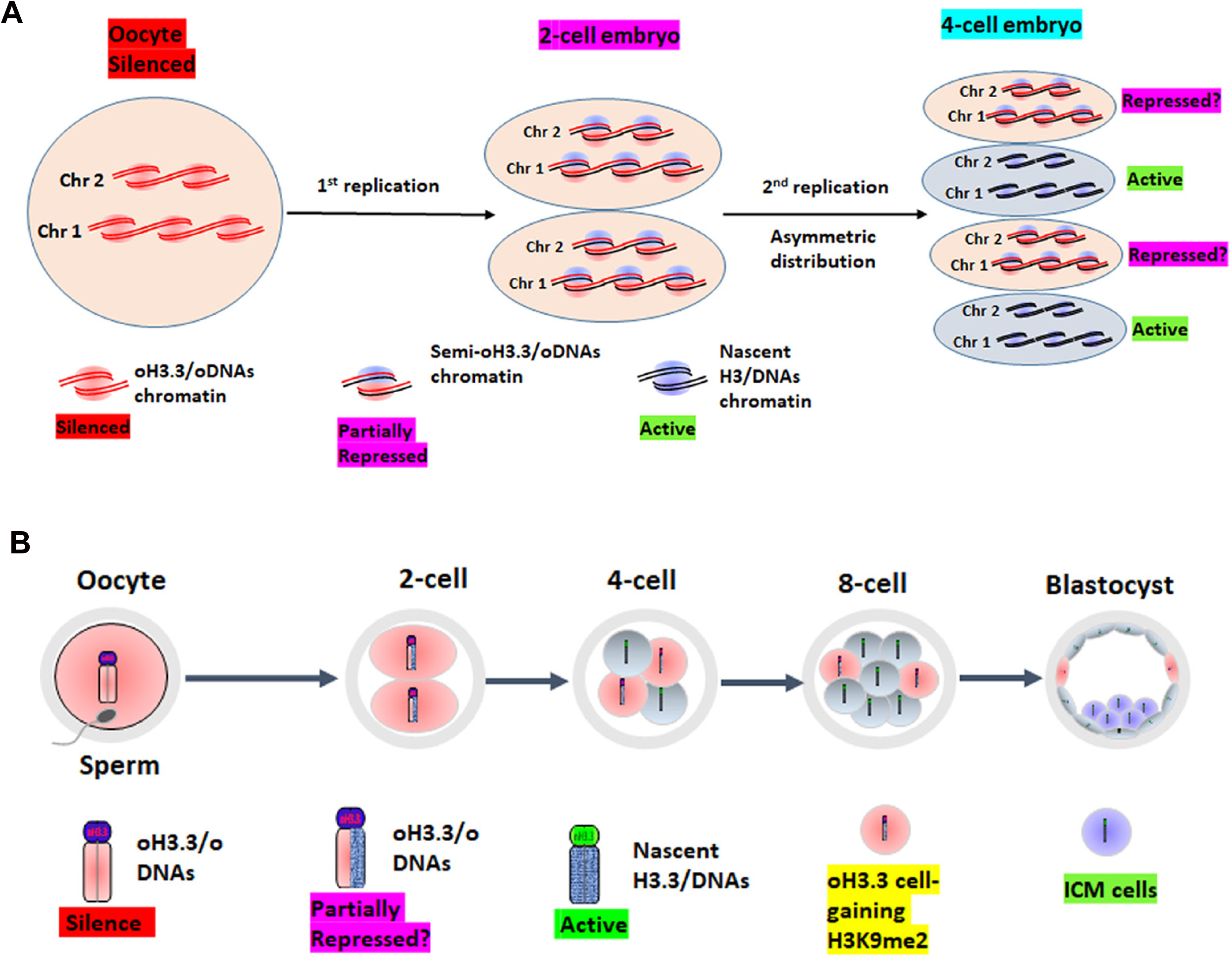
A model of oH3.3-mediated maternal genome activation during embryogenesis. (**A**) Schematic illustration of oH3.3/oDNA segregation and distribution during the first and second embryonic divisions. The oH3.3-associated chromatin remains in a repressed state. The oH3.3 is symmetrically distributed between the two daughter chromatids, which each double-stranded DNA consists of one nascent DNA strand (represented by a black line) and one old strand from the parental oocyte DNA (oDNAs, represented by a red line) during the first round of DNA replication. After the second round of DNA replication, the segregation and distribution of oH3.3 take an asymmetric pattern. Chromatids containing oH3.3/oDNA are allocated together into one cell, generating two cells with retained oH3.3/oDNA and two cells with nascent H3/DNAs at the four-cell stage. The cells retaining oH3.3/oDNAs remain partially repressed and continue with asymmetric distribution in the following DNA replications. The maternal genome is fully activated in the cells with nascent H3/DNAs, while the maternal genome in the two cells retaining oH3.3/oDNAs remains partially repressed. Chr 1 and Chr 2 serve as examples of the 20 maternal chromosomes in mice. (**B**) In preimplantation embryos, chromosomes associated with oH3.3 co-segregate. One strand of each chromosome from the oocyte genome is retained in one cell, while the counterpart strand is inherited by another cell during mitosis. This asymmetric distribution of oH3.3/oDNAs is facilitated by a dual-spindle formation mechanism. Cells that retain oH3.3/oDNA acquire the constitutive heterochromatin marker H3K9me2 and do not contribute to the inner cell mass (ICM) of a blastocyst. Consequently, these cells are excluded from the developing embryo.

## Discussion

Our research in mice has revealed a remarkable phenomenon in early embryonic development – the asymmetric segregation and distribution of the H3.3 protein associated with the oocyte genome (oH3.3). During preimplantation, oH3.3 is selectively retained in only two blastomeres within fertilized embryos. These oH3.3-retaining cells exhibit distinct characteristics, including a more compact nucleus and positive staining for the constitutive heterochromatin marker, H3K9me2. These blastomeres preferentially contribute to the trophoblast layer rather than to the inner cell mass (ICM) in the blastocyst, and a majority of blastocysts lose one oH3.3-retained cell. Furthermore, we have observed that the full activation of the maternal genome is delayed until the 4-cell stage. As oH3.3 is associated with oocyte DNA, it suggests the co-segregation and distribution of oocyte DNA strands (oDNAs): One strand of each chromosome of the oocyte genome is retained in one cell, while the other strand is taken away by another cell during early embryogenesis. This delay in maternal genome activation may be due to repressive epigenetic modifications in oocyte DNAs that persist in the zygote and the 2-cell stage embryo (7, 11, 12). In contrast, the paternal genome undergoes active demethylation (8, 12, 33). This delay aligns with a higher transcriptional activity in the paternal genome compared to the maternal genome within zygotes (13). It also corresponds to observations in humans where paternal genome activation precedes maternal genome activation (14).

Our proposed model suggests that oH3.3-associated oDNAs remain partially repressed due to the persistence of oocyte-specific repressive epigenetic modifications. Full activation of the maternal genome requires DNA replication to generate nascent maternal DNAs free of these repressive modifications. This occurs after the second DNA replication and in the four-cell stage embryo (**Fig. 6A**). Mechanistically, during embryonic DNA replication, oH3.3 is initially symmetrically distributed between the two daughter chromatids. However, after the second round of DNA replication, asymmetric segregation occurs. oH3.3 is asymmetrically segregated with parental oDNAs, while the nascent histone H3 is distributed to newly synthesized DNAs. All twenty chromatids containing oH3.3/oDNAs are allocated into a single cell at the four-cell stage, breaking the symmetry. Two blastomeres retain all oH3.3/oDNAs chromatids, remaining repressed, while the other two receive nascent histone H3/DNAs chromatids, allowing full activation at the four-cell stage (**Fig. 6A**).

Our model outlines two crucial mechanisms for maternal genome activation: asymmetric nucleosome segregation of oH3.3 and the distribution of all oocyte chromatids into one cell. This challenges the conventional view of H3.3 deposition and highlights oH3.3’s role in the global silencing of the oocyte genome. Although asymmetrical inheritance of histones is known in other species (34, 35), understanding in mammals has been limited. Repressed chromatin transmission during cell division is critical to maintaining cell identity (36–39). Our findings provide a model for this process involving a globally repressed genome at the cellular level in mammals. In mammals, parental chromosomes initially converge during zygotic mitosis. Recent research reveals separate spatial domains for parental genomes during the first mitosis (40, 41), aiding the segregation and distribution of oH3.3-containing sister chromatids from the maternal genome into one cell. This process separates oH3.3-containing chromatids with oocyte-specific modifications, facilitating full maternal genome activation. This critical mechanism for embryonic development was previously unrecognized, a highlight of our study.

Although the random distribution of sperm DNA among blastomeres is known (42), observing the segregation and distribution of oocyte DNA (oDNAs) directly presents technical challenges. Our research strongly suggests asymmetric oDNA distribution, although concrete evidence is currently lacking. During blastocyst formation, cells with oH3.3/oDNAs favor trophoblast differentiation over epiblasts. This bias aligns with observed gene expression patterns (43, 44). This strategy may minimize transcriptional errors such as maternal imprinting genes, ensuring robust post-implantation embryonic development (**Fig.6B**).

In summary, our discovery of oH3.3’s asymmetric distribution sheds light on the complex processes governing epigenetic symmetry breaking and maternal genome activation in early embryonic development. More research is needed to understand the precise roles of oH3.3 and oDNAs and whether similar mechanisms exist in other species.

## Acknowledgments

I extend my gratitude to Drs. Danny Reinberg, Xin Chen, Jose Silva, Jianlong Wang, Wei Xie, Fengling Chen, Qing-Yuan Sun, and the members of my laboratory for their insightful discussions and invaluable suggestions. D.W. was supported by NIH grants (GM129380 and 1R21OD031973) and New York State Stem Cell Science Program (NYSTEM; contract C32581GG). This study was partially supported by Starr Foundation Stem Cell Core Project and initiatives TRI-SCI 2019-029 (to Z.R.).

## Author contributions

D.W. conceived the project, performed the experiments, and wrote the paper. Z.R. provide financial support and offered insightful discussions.

## Declaration of interests

Authors declare that they have no competing interests.

## Methods

### Animals and oocytes

Animals were housed and prepared following the protocol approved by the IACUC of Weill Cornell Medical College (Protocol number: 2014-0061). B6D2F1 and ICR mice were obtained from Taconic Farms (Germantown, NY). Female mice, aged 6-8 weeks, were superovulated using 5 IU of PMSG (Pregnant mare serum gonadotrophin, Sigma-Aldrich, St. Louis, MO) and 5 IU of hCG (Human chorionic gonadotrophin, Sigma-Aldrich) with a 48-hour interval between injections. MII oocytes were collected from superovulated female mice 14-16 hours after hCG administration.

### Injection of siRNA into MII oocytes

MII oocytes were collected from superovulated B6D2F1 females 14-16 hours after hCG injection. siRNAs were introduced into the oocytes using a piezo-operated microcapillary pipette with a 3-5 µm inner diameter. Post-injection, oocytes were allowed to rest at room temperature for 10 minutes before being transferred to an incubator for at least an additional 30 minutes. siRNA sequences for H3.3A (*h3f3a*) and H3.3B (*h3f3b*) as following(19):

*h3f3a* 1s – CGUUCAUUUGUGUGUGAAUUUUU;

*h3f3a* 1as – AAAUUCACACACAAAUGAACGUU;

*h3f3a* 2s – GCGAGAAAUUGCUCAGGACUUUU;

*h3f3a* 2as – AAGUCCUGAGCAAUUUCUCGCUU;

*h3f3b* 1s – UCUGAGAGAGAUCCGUCGUUAUU;

*h3f3b* 1as – UAACGACGGAUCUCUCUCAGAUU;

*h3f3b* 2s – GAAGCUGCCAUUCCAGAGAUUUU;

*h3f3b* 2as – AAUCUCUGGAAUGGCAGCUUCUU

### Spindle Exchange, Nuclear transfer, and Intracytoplasmic Sperm Injection (ICSI)

Oocytes (both H3.3B-HA and B6D2F1) were transferred to a droplet of HEPES-CZB medium containing 5 µg/ml cytochalasin B, and preplaced in the operation chamber on the microscope stage. Using a micromanipulator and a fine glass needle, the spindle-chromosomal complex (SCC) was carefully removed from both donor H3.3B-HA oocytes and recipient B6D2F1 oocytes. The SCC from the H3.3B-HA oocyte was then transferred to the perivitelline space of the enucleated B6D2F1 oocyte. The oocytes subsequently underwent electrofusion, during which they were aligned, and a series of electrical pulses were applied to induce membrane fusion. The fused oocytes were activated by culturing in Ca^2+^-free CZB medium containing both 10 mM Sr2+ and 5 µg/ml cytochalasin B for 5 hours. The oocytes were then transferred to an incubator and cultured in advanced KSOM (Cat # MR-101-D, Millipore) at 37°C under 5% (v/v) CO2 in air.

Nuclear transfer was performed following established protocols by Wakayama et al., Nature, 1998. Oocyte enucleation was carried out as described earlier. Post enucleation, oocytes were transferred to cytochalasin B-free KSOM and placed back in the incubator. The donor nuclei (Oct4-EGFP ESCs) were gently aspirated in and out of the injection pipette until their nuclei were largely devoid of visible cytoplasmic material. Each nucleus was then injected into a separate enucleated oocyte.

After somatic-cell nucleus injection, oocytes were activated by culturing in Ca2+-free CZB containing both 10 mM Sr2+ and 5 µg ml-1 cytochalasin B for 5 hours. Subsequently, they were further cultured in the medium supplemented with aphidicolin (5 μg/ml) to inhibit DNA synthesis for 3 days, which corresponds to the morula stage. The incubation was carried out at 37°C under 5% (v/v) CO2 in air.

For ICSI, a WT sperm head from ICR mice was picked up in the PVP droplet using the injection pipette, and each H3.3B-HA oocyte was injected with one sperm head. Following ICSI, oocytes were allowed to rest at room temperature for 10 minutes before being transferred to KSOM medium for culture in the incubator at 37°C under 5% (v/v) CO2 in air.

### Fluorescence microscopy, immunohistochemistry, and confocal imaging

Fluorescence expression was detected in live embryos using a fluorescence inverted microscope (Nikon, TE2000-U). Images were captured with a digital camera and merged using NIS-Elements D software (Nikon). For immunohistochemistry staining (IF), oocytes or embryos were fixed in 4% paraformaldehyde, permeabilized with 0.5% Triton X-100 in PBS, and blocked using a solution containing 10% normal donkey serum and 0.5% Triton in PBS. Oocytes or embryos were then incubated in working dilutions of the primary antibodies, including anti-HA goat IgG (Abcam, ab9134, 1:100) and anti-α-tubulin-FITC (Sigma, F2168, 1:300), Anti-Histone H3 (di methyl K9) (H3K9me2) (Abcam, ab1220, 1:100). As a secondary antibody, anti-goat IgG conjugated with Alexa Fluor 647 (Invitrogen, A-21245) was used. Imaging was performed using a Zeiss 710 confocal imaging system. Z-stack images of 20 sequential sections for each embryo were captured.

## Notes

### Competing Interest Statement

The authors have declared no competing interest.

## References

1. Smallwood, S. A., Tomizawa, S., Krueger, F., Ruf, N., Carli, N., Segonds-Pichon, A., Sato, S., Hata, K., Andrews, S. R., and Kelsey, G. (2011) Dynamic CpG island methylation landscape in oocytes and preimplantation embryos. Nat Genet 43, 811–U126

2. Shirane, K., Toh, H., Kobayashi, H., Miura, F., Chiba, H., Ito, T., Kono, T., and Sasaki, H. (2013) Mouse Oocyte Methylomes at Base Resolution Reveal Genome-Wide Accumulation of Non-CpG Methylation and Role of DNA Methyltransferases. Plos Genet 9

3. Guo, H. S., Zhu, P., Yan, L. Y., Li, R., Hu, B. Q., Lian, Y., Yan, J., Ren, X. L., Lin, S. L., Li, J. S., Jin, X. H., Shi, X. D., Liu, P., Wang, X. Y., Wang, W., Wei, Y., Li, X. L., Guo, F., Wu, X. L., Fan, X. Y., Yong, J., Wen, L., Xie, S. X., Tang, F. C., and Qiao, J. (2014) The DNA methylation landscape of human early embryos. Nature 511, 606-+

4. Kobayashi, H., Sakurai, T., Imai, M., Takahashi, N., Fukuda, A., Yayoi, O., Sato, S., Nakabayashi, K., Hata, K., Sotomaru, Y., Suzuki, Y., and Kono, T. (2012) Contribution of Intragenic DNA Methylation in Mouse Gametic DNA Methylomes to Establish Oocyte-Specific Heritable Marks. Plos Genet 8

5. Ishiuchi, T., Abe, S., Inoue, K., Yeung, W. K. A., Miki, Y., Ogura, A., and Sasaki, H. (2021) Reprogramming of the histone H3.3 landscape in the early mouse embryo. Nat Struct Mol Biol 28

6. Lee, M. T., Bonneau, A. R., and Giraldez, A. J. (2014) Zygotic Genome Activation During the Maternal-to-Zygotic Transition. Annu Rev Cell Dev Bi 30, 581–613

7. Rougier, N., Bourc’his, D., Gomes, D. M., Niveleau, A., Plachot, M., Paldi, A., and Viegas-Pequignot, E. (1998) Chromosome methylation patterns during mammalian preimplantation development. Genes & development 12, 2108–2113

8. Gu, T. P., Guo, F., Yang, H., Wu, H. P., Xu, G. F., Liu, W., Xie, Z. G., Shi, L. Y., He, X. Y., Jin, S. G., Iqbal, K., Shi, Y. J. G., Deng, Z. X., Szabo, P. E., Pfeifer, G. P., Li, J. S., and Xu, G. L. (2011) The role of Tet3 DNA dioxygenase in epigenetic reprogramming by oocytes. Nature 477, 606–U136

9. Smith, Z. D., Chan, M. M., Humm, K. C., Karnik, R., Mekhoubad, S., Regev, A., Eggan, K., and Meissner, A. (2014) DNA methylation dynamics of the human preimplantation embryo. Nature 511, 611–+

10. Wu, X. J., and Zhang, Y. (2017) TET-mediated active DNA demethylation: mechanism, function and beyond. Nat Rev Genet 18, 517–534

11. Jenkins, T. G., and Carrell, D. T. (2012) Dynamic alterations in the paternal epigenetic landscape following fertilization. Front Genet 3, 143

12. Mayer, W., Niveleau, A., Walter, J., Fundele, R., and Haaf, T. (2000) Embryogenesis - Demethylation of the zygotic paternal genome. Nature 403, 501–502

13. Aoki, F., Worrad, D. M., and Schultz, R. M. (1997) Regulation of transcriptional activity during the first and second cell cycles in the preimplantation mouse embryo. Developmental biology 181, 296–307

14. Yuan, S. L., Zhan, J. H., Zhang, J. Y., Liu, Z. B., Hou, Z. Z., Zhang, C. X., Yi, L. Z., Gao, L., Zhao, H., Chen, Z. J., Liu, J., and Wu, K. L. (2023) Human zygotic genome activation is initiated from paternal genome. Cell Discov 9

15. Reichmann, J., Nijmeijer, B., Hossain, M. J., Eguren, M., Schneider, I., Politi, A. Z., Roberti, M. J., Hufnagel, L., Hiiragi, T., and Ellenberg, J. (2018) Dual-spindle formation in zygotes keeps parental genomes apart in early mammalian embryos. Science 361, 189–193

16. Frank, D., Doenecke, D., and Albig, W. (2003) Differential expression of human replacement and cell cycle dependent H3 histone genes. Gene 312, 135–143

17. Wellman, S. E., Casano, P. J., Pilch, D. R., Marzluff, W. F., and Sittman, D. B. (1987) Characterization of mouse H3.3-like histone genes. Gene 59, 29–39

18. Szenker, E., Ray-Gallet, D., and Almouzni, G. (2011) The double face of the histone variant H3.3. Cell Research 21, 421–434

19. Wen, D. C., Banaszynski, L. A., Liu, Y., Geng, F. Q., Noh, K. M., Xiang, J., Elemento, O., Rosenwaks, Z., Allis, C. D., and Rafii, S. (2014) Histone variant H3.3 is an essential maternal factor for oocyte reprogramming. Proceedings of the National Academy of Sciences of the United States of America 111, 7325–7330

20. Guo, P. P., Liu, Y., Geng, F. Q., Daman, A. W., Liu, X. Y., Zhong, L. W., Ravishankar, A., Lis, R., Duran, J. G. B., Itkin, T., Tang, F. Y., Zhang, T., Xiang, J., Shido, K., Ding, B. S., Wen, D. C., Josefowicz, S. Z., and Rafii, S. (2022) Histone variant H3.3 maintains adult haematopoietic stem cell homeostasis by enforcing chromatin adaptability (vol 24, pg 99, 2022). Nat Cell Biol 24, 279–279

21. Jang, C. W., Shibata, Y., Starmer, J., Yee, D., and Magnuson, T. (2015) Histone H3.3 maintains genome integrity during mammalian development. Genes & development 29, 1377–1392

22. Wen, D., Noh, K. M., Goldberg, A. D., Allis, C. D., Rosenwaks, Z., Rafii, S., and Banaszynski, L. A. (2014) Genome editing a mouse locus encoding a variant histone, H3.3B, to report on its expression in live animals. Genesis 52, 959–966

23. Kong, Q. R., Banaszynski, L. A., Geng, F. Q., Zhang, X. L., Zhang, J. M., Zhang, H., O’Neill, C. L., Yan, P. D., Liu, Z. H., Shido, K., Palermo, G. D., Allis, C. D., Rafii, S., Rosenwaks, Z., and Wen, D. C. (2018) Histone variant H3.3-mediated chromatin remodeling is essential for paternal genome activation in mouse preimplantation embryos. Journal of Biological Chemistry 293, 3829–3838

24. Lin, C. J., Conti, M., and Ramalho-Santos, M. (2013) Histone variant H3.3 maintains a decondensed chromatin state essential for mouse preimplantation development. Development 140, 3624–3634

25. Padeken, J., Methot, S. P., and Gasser, S. M. (2022) Establishment of H3K9-methylated heterochromatin and its functions in tissue differentiation and maintenance. Nat Rev Mol Cell Bio 23, 623–640

26. Yeung, W. K. A., Brind’Amour, J., Hatano, Y., Yamagata, K., Feil, R., Lorincz, M. C., Tachibana, M., Shinkai, Y., and Sasaki, H. (2019) Histone H3K9 Methyltransferase G9a in Oocytes Is Essential for Preimplantation Development but Dispensable for CG Methylation Protection. Cell Rep 27, 282-+

27. Wen, B., Wu, H., Shinkai, Y., Irizarry, R. A., and Feinberg, A. P. (2009) Large histone H3 lysine 9 dimethylated chromatin blocks distinguish differentiated from embryonic stem cells. Nat Genet 41, 246–250

28. Cheloufi, S., Elling, U., Hopfgartner, B., Jung, Y. L., Murn, J., Ninova, M., Hubmann, M., Badeaux, A. I., Ang, C. E., Tenen, D., Wesche, D. J., Abazova, N., Hogue, M., Tasdemir, N., Brumbaugh, J., Rathert, P., Jude, J., Ferrari, F., Blanco, A., Fellner, M., Wenzel, D., Zinner, M., Vidal, S. E., Bell, O., Stadtfeld, M., Chang, H. Y., Almouzni, G., Lowe, S. W., Rinn, J., Wernig, M., Aravin, A., Shi, Y., Park, P. J., Penninger, J. M., Zuber, J., and Hochedlinger, K. (2015) The histone chaperone CAF-1 safeguards somatic cell identity. Nature 528, 218–+

29. Djekidel, M. N., Inoue, A., Matoba, S., Suzuki, T., Zhang, C. X., Lu, F. L., Jiang, L., and Zhang, Y. (2018) Reprogramming of Chromatin Accessibility in Somatic Cell Nuclear Transfer Is DNA Replication Independent. Cell Rep 23, 1939–1947

30. Jullien, J., Astrand, C., Halley-Stott, R. P., Garrett, N., and Gurdon, J. B. (2010) Characterization of somatic cell nuclear reprogramming by oocytes in which a linker histone is required for pluripotency gene reactivation. Proceedings of the National Academy of Sciences of the United States of America 107, 5483–5488

31. Tomikawa, J., Penfold, C. A., Kamiya, T., Hibino, R., Kosaka, A., Anzai, M., Matsumoto, K., and Miyamoto, K. (2021) Cell division-and DNA replication-free reprogramming of somatic nuclei for embryonic transcription. Iscience 24

32. Deng, Q. L., Ramskold, D., Reinius, B., and Sandberg, R. (2014) Single-Cell RNA-Seq Reveals Dynamic, Random Monoallelic Gene Expression in Mammalian Cells. Science 343, 193–196

33. Oswald, J., Engemann, S., Lane, N., Mayer, W., Olek, A., Fundele, R., Dean, W., Reik, W., and Walter, J. (2000) Active demethylation of the paternal genome in the mouse zygote. Current Biology 10, 475–478

34. Wooten, M., Snedeker, J., Nizami, Z. F., Yang, X., Ranjan, R., Urban, E., Kim, J. M., Gall, J., Xiao, J., and Chen, X. (2019) Asymmetric histone inheritance via strand-specific incorporation and biased replication fork movement. Nat Struct Mol Biol 26, 732–743

35. Urban, J. A., Ranjan, R., and Chen, X. (2022) Asymmetric Histone Inheritance: Establishment, Recognition, and Execution. Annu Rev Genet 56, 113–143

36. Reinberg, D., and Vales, L. D. (2018) Chromatin domains rich in inheritance. Science 361, 33–34

37. Saxton, D. S., and Rine, J. (2019) Epigenetic memory independent of symmetric histone inheritance. Elife 8

38. Escobar, T. M., Oksuz, O., Saldana-Meyer, R., Descostes, N., Bonasio, R., and Reinberg, D. (2019) Active and Repressed Chromatin Domains Exhibit Distinct Nucleosome Segregation during DNA Replication. Cell 179, 953-+

39. Escobar, T. M., Yu, J. R., Liu, S., Lucero, K., Vasilyev, N., Nudler, E., and Reinberg, D. (2022) Inheritance of repressed chromatin domains during S phase requires the histone chaperone NPM1. Sci Adv 8, eabm3945

40. Mayer, W., Smith, A., Fundele, R., and Haaf, T. (2000) Spatial separation of parental genomes in preimplantation mouse embryos. J Cell Biol 148, 629–634

41. van de Werken, C., van der Heijden, G. W., Eleveld, C., Teeuwssen, M., Albert, M., Baarends, W. M., Laven, J. S. E., Peters, A. H. F. M., and Baart, E. B. (2014) Paternal heterochromatin formation in human embryos is H3K9/HP1 directed and primed by sperm-derived histone modifications. Nat Commun 5

42. Ito, K., Mcghee, J. D., and Schultz, G. A. (1988) Paternal DNA Strands Segregate to Both Trophectoderm and Inner Cell Mass of the Developing Mouse Embryo. Genes & development 2, 929–936

43. Miyanari, Y., and Torres-Padilla, M. E. (2012) Control of ground-state pluripotency by allelic regulation of Nanog. Nature 483, 470–473

44. Silva, J., Nichols, J., Theunissen, T. W., Guo, G., van Oosten, A. L., Barrandon, O., Wray, J., Yamanaka, S., Chambers, I., and Smith, A. (2009) Nanog Is the Gateway to the Pluripotent Ground State. Cell 138, 722–737

